# Gene Expression Plasticity Is Associated with Regulatory Complexity but Not with Specific Network Motifs

**DOI:** 10.1101/2024.03.11.584403

**Authors:** Apolline J. R. Petit, Anne Genissel, Arnaud Le Rouzic

**Affiliations:** Université Paris-Saclay, CNRS, IRD, UMR EGCE, 91190, Gif-sur-Yvette, France; Université Paris Saclay, INRAE, UR BIOGER, 91120, Palaiseau, France

**Author notes:** Corresponding author: Université Paris-Saclay, CNRS, IRD, UMR EGCE, 91190, Gif-sur-Yvette, France.

## Abstract

Over the past two decades, a large body of theoretical and empirical work has been conducted with the aim of identifying the gene regulatory network topologies responsible for gene expression dynamics. Some studies have linked gene expression plasticity to specific network motifs such as feedforward loops, diamond motifs, or feedback loops. However, both theoretical and empirical work have produced equivocal results, as the same topologies have also been associated with expression robustness. As a step toward understanding the regulatory basis of gene expression plasticity, our goal is to understand how local regulatory topologies may contribute to it. To this end we compared theoretical predictions from a simulated network evolution model with empirical data based on *Escherichia coli* regulatory network. We investigated the link between network topology and gene expression at three levels: the number of regulators, the number and the proportion of loops (feedback loops, feedforward loops and diamond loops), and the proportion of unique motifs (here characterized by the position of up- or down-regulations within loops). Consistent results from our empirical and theoretical approaches revealed that plastic genes are, on average, regulated by a greater number of genes. In addition, our theoretical predictions showed that selection, as opposed to genetic drift, strongly biases the distribution of network motifs. However, similar results were observed when comparing plastic and non-plastic genes. Overall, this work illustrates that our current understanding of network topology may be insufficient to fully explain or predict gene expression plasticity.

## Introduction

Understanding the mechanisms involved in the evolution of regulatory networks remains a major challenge in evolutionary and systems biology. It is widely acknowledged that biological gene regulatory networks are not randomly organized (Shen-Orr *et al.*, 2002; Burda *et al.*, 2011), but the interplay between natural selection, mutation bias, functional constraints, and genetic drift in the evolution of network topology is not clearly established. Some network topologies, such as feedforward loops (FFLs) and feedback loops (FBLs) have attracted considerable attention because of their abundance in biological systems (Mangan and Alon, 2003; Alon, 2007). This enrichment could be the consequence of selection indirectly promoting motifs through their effect on gene expression dynamics, including stability, plasticity, or oscillation.

Changes in gene expression is a molecular manifestation of phenotypic plasticity and it conditions the tempo and mode of the organism’s response to environmental cues. At the gene level, plasticity is contingent on signal-sensitive regulation of transcription, and selection for plasticity might indirectly affect the underlying regulatory network. For instance, Vlková and Silander, 2022 demonstrated that *Escherichia coli* promoters involved in the metabolism of specific substrate are often selected to maximize plasticity. Yet, both theoretical and empirical work falls short of consensus when linking gene expression dynamics with specific network topology. From the theoretical side, feedforward loops (FFLs) (Shen-Orr *et al.*, 2002; Basu *et al.*, 2004; Ma *et al.*, 2009; de Ronde *et al.*, 2012; Xiong *et al.*, 2019; Wang *et al.*, 2020), diamond (DMD) loops (de Ronde *et al.*, 2012; Xiong *et al.*, 2019), and feed-back loops (FBLs) (Hornung and Barkai, 2008; Panovska-Griffiths *et al.*, 2013) have been reported to be involved in plastic (environmentally-sensitive) gene expression. Counterintuitively, other theoretical findings associate those same loop categories with robustness-related functions (FFL: Osella *et al.*, 2011; Kadelka *et al.*, 2013; Roy *et al.*, 2017, DMD: Khammash, 2021, FBL: Kwon and Cho, 2007; Wang *et al.*, 2020). From the empirical side, many studies observed these motifs in both plastic and robust systems. Two studies on synthetic bacteria (Basu *et al.*, 2004) and *E. coli* (Sen *et al.*, 2014) brought experimental evidence of a link between feedforward loops and sensitivity to environmental signals. However these studies were restricted to a few genes, and the genome-wide picture remains unknown (Frei and Khammash, 2021; Weldemichael *et al.*, 2022). Overall, there is no clear indication whether specific network topologies are involved in the regulation of gene expression plasticity.

A considerable obstacle in the quest to identify biologically relevant topologies in networks is the lack of a clear theoretical framework indicating where information should be sought. Gene network topology can be described by multiple indicators that focus on different scales, including the entire network, the gene level (e.g., the number of incoming or outgoing connections), and local regulatory topologies, such as the nature of regulatory loops (feedback, feedforward, or diamond) and the loop motifs (the arrangement of up- and down-regulation within loops). Which of these levels, if any, is relevant for gene expression plasticity, and is it possible to explain the contradictory results from the literature? Our ambition here is to study the effect of local regulatory topologies in networks with a dual approach, coupling a theoretical model with a data-driven regulatory network analysis, in order to identify consistent predictions and observations. We derived theoretical predictions from evolutionary simulations, in which the regulatory interactions were free to evolve under a mutation-drift-selection process (need a reference here for this model?). Gene expression plasticity was then estimated as the response of individual genes to an environmental signal. In parallel, we collected lists of genes significantly responding to various environmental conditions according to 11 transcriptomic studies in *E. coli*. Despite that many regulatory pathways are known for this model species, the link between local regulatory network topology and gene expression plasticity is poorly investigated. In this study, we made use of one of the most up-to-date transcriptional regulation databases, EcoCyc (Keseler *et al.*, 2021), to retrieve regulatory interactions. Both simulated and empirical datasets were then analyzed within the same statistical framework, enabling direct comparison. We found that genes exhibiting expression plasticity were regulated by more genes on average. In addition, this study showed that selection biased the distribution of loop motifs, regardless of the level of gene expression plasticity.

## Material and Methods

### Empirical data: *E. coli*

We used *E. coli* K-12 substr. MG1655 annotation (RefSeq: GCF_000005845.2). *E. coli* K-12 known regulations have been obtained from Ecocyc v.28.5 (Keseler *et al.*, 2021, downloaded 29/01/2024). This release contained 9534 regulatory interactions.

To conduct our analyses, we established a list of “plastic” genes, to be contrasted to “non-plastic” genes. We defined plasticity as gene expression that respond to several environmental cues, with strong enough amplitude that it could be detected and statistically significant. Non-plasticity was attributed to genes never differentially expressed in any environmental cues considered. We selected 11 studies explicitly testing environmental conditions on gene expression in *E. coli*, and proposing lists of differentially-expressed genes. The environmental factors were pH (Tucker *et al.*, 2002; Maurer *et al.*, 2005), growth medium (urine, gluconate, and lysogeny broth) (Feugeas *et al.*, 2016), aerosolization and liquid suspension (Ng *et al.*, 2018), exposure to apple juice (Bergholz *et al.*, 2009), magnesium stress, sodium stress and carbon source (Caglar *et al.*, 2017), temperature (White-Ziegler *et al.*, 2007; Kim *et al.*, 2020), metabolic stress (low pH, low oxygen, low nutrients) (Bhatia *et al.*, 2022), oxidative stress (Wang *et al.*, 2009), and amino acid starvation (Durfee *et al.*, 2008). Genes were considered as plastic when reported as differentially expressed in at least two independent studies. By contrast genes were considered as “non-plastic” if they were never reported as differentially expressed in these studies.

### Gene regulatory network simulations

We used a quantitative gene regulatory network model, inspired by A. Wagner’s seminar work on gene expression robustness and plasticity (Wagner, 1994, 1996). This setting, coupling a traditional population genetics model with a gene network model as a genotype-to-phenotype map, was particularly popular to explore the evolution of complex genetic architectures (Draghi and Wagner, 2008; Ng and Kinjo, 2023; Petit *et al.*, 2023).

**Gene network** The gene network model was dynamic with discrete time-steps. Genotype consisted in *n* = 36 genes encoding a gene network represented by a *n* ×*n* gene regulatory matrix **W**. Genes were expressed quantitatively between *P* = 0 (no expression) and *P* = 1 (maximum expression). Initial gene expressions were set to their basal level (constitutive expression) *κ* = 0.15, representing the gene expression in absence of regulation, and updated as a function of the concentration of the other genes in the network during 24 time steps:

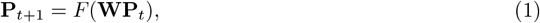

where **P** was a vector of *n* quantitative gene expressions scaled between 0 and 1 by a multivariate sigmoid function *F* (*x*_1_, *…, x*_*n*_) = (*f* (*x*_1_), *…, f* (*x*_*n*_)), with *f* (*x*) = [1 + (1*/κ −*1) exp(−*x/κ*(1 −*κ*))]^*−1*^ as in Rünneburger and Le Rouzic, 2016.

Individuals were characterized by their phenotype, consisting in the average expression of their *n* genes during the four last time-steps 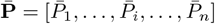, with 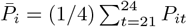.

### Reproduction, mutation, and selection

Populations consisted in 20,000 haploid, clonally-reproducing individuals, parameterized to match approximately *E. coli* ‘s life cycle. Simulations were carried on for 4,000 non-overlapping generations.

The genotype **W** was affected by mutations of the regulation strength *W*_*ij*_, at a rate of *µ* = 0.15 per individual. Each gene had the same probability *µ/n* = 0.004 to be affected by a mutation. A mutated gene *i* had a random regulation *W*_*ij*_ shifted by a Gaussian random number of standard deviation 0.05. This setting (many mutations of small effect) ensured a substantial amount of polymorphism in the population, so that evolution was not limited by mutations.

Successive generations were produced by drawing clones with a probability proportional to their relative fitness. The individual fitness *w* was the product of two components: *w* = *w*_*e*_ × *w*_*s*_. The first term *w*_*e*_ was determined by a stabilizing bell-shaped fitness function and decreased with the Euclidean distance to the gene expression optima ***θ*** (Lande, 1979):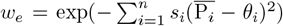. Unless specified otherwise, the selection coefficient was *s*_*i*_ = 0 for “unselected” genes, and *s*_*i*_ = 10 for “selected” genes. The second term *w*_*s*_ corresponds to the selection for network stability, calculated as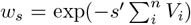, where *s*^*′*^ quantifies the selection against unstable gene expression, and 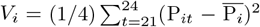 was the variance in the expression of gene *i* during the last 4 steps of the network dynamics. We set *s*^*′*^ = 46, 000 (as in Rünneburger and Le Rouzic, 2016; Petit *et al.*, 2023), which was a large penalty, reflecting the popular conjecture that unstable networks are not viable (Siegal and Bergman, 2002).

### Fluctuating selection and plasticity

In our simulations, the environment was modeled as a numerical value *E*_*g*_ changing randomly every generation *g*, drawn from an uniform distribution *U* (0.15, 0.85). The state of the environment was transmitted to the network via a sensor gene, as in Draghi and Whitlock, 2012; Vadée-Le-Brun *et al.*, 2015. The model implementation follows Burban *et al.*, 2022, the expression of the sensor gene was constant *P*_*s*_ = *E*_*g*_ during all the network dynamics. The sensor gene was free to evolve regulations toward any genes of the network, but it was not genetically regulated, as its expression depends solely of the environment.

The other 35 network genes were divided in 4 categories, see Table 1. 10 genes were “Transcription factors” not subjected to direct selection; 10 “Plastic Target Genes” were directly selected to have their expression correlated with the environment; 10 “Non-plastic Target Genes” were directly selected to have a constant expression; and 5 unselected “neutral genes” were let free to evolve.

**Table 1:** Description of the five gene categories implemented in the gene regulatory network model.

|  | Sensor gene | Transcription factor (TF) | Plastic Target gene | Constant Target gene | Neutral gene |
| --- | --- | --- | --- | --- | --- |
| Number | 1 | 10 | 10 | 10 | 5 |
| Selection $s_i$ | 0 | 0 | 10 | 10 | 0 |
| Optimum $\theta_i$ | | | $a_i E_g + b_i$ | $b_i$ | |
| Evolution | Environment only | Free | Towards plasticity | Towards robustness | Drift |
| Regulating | All genes | All genes but sensor | TF | TF | None |
| Regulated by | None | All genes but Neutral | TF and Sensor | TF and Sensor | TF and Sensor |

Selection on plastic genes was modelled by changing the gene expression optimum every generation *g* as a function of the environment *E*_*g*_, as *θ*_*i*_ = *a*_*i*_*E*_*g*_ + *b*_*i*_. The intercept *b*_*i*_ and the slope *a*_*i*_ were randomly drawn for each gene *i* at the beginning of each simulation under the constraint that optimal gene expression had to remain between 0 and 1 for any *E*_*g*_ ∈ (0.15, 0.85), and stayed constant over generations. “Non-plastic” genes had their constant optimum drawn from a uniform distribution *U* (0.15, 0.85).

The reaction norm is the function describing the trait response to an environmental variation. We measured the expression plasticity of a gene *i* as the slope *β*_*i*_ of the linear regression between a set of environments and the corresponding gene expression 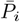. Simulated genes were classified into the category “plastic genes” and “non-plastic genes” according to their reaction norm slope. Genes with *|β_i_|* ≥ 0.5 were considered as plastic and genes with *|β_i_|* ≤ 0.025 as non-plastic. Genes outside of these categories were not kept during the analysis.

### Network Analysis

We used *E. coli* Ecocyc regulation data to construct a discrete 4638-gene network as a 4638× 4638 square regulatory matrix **Ec**. A cell Ec_*ij*_ ∈{−1, 0, 1} represents the regulation of gene *j* on *i*. Regulations that were both activating and inhibiting their target in the Ecocyc dataset were considered as null.

To study the regulatory motifs responsible for plasticity in the simulated networks with the same pipeline used for *E. coli*, we discretized the regulation matrix **W** into a matrix **Et**, based on the concept of “essentiality” (Burda *et al.*, 2011). We tested one by one every element of the **W** matrix by changing their value to 0. If it changed the reaction norm slope or the intercept by more than 0.005, then the regulation was considered as “essential”: Et_*ij*_ = sign(W_*ij*_), and Et_*ij*_ = 0 otherwise.

#### Motifs

We investigated three loop categories: 3-gene feedforward loops (corresponding to 8 unique motifs, Table 2), four-gene diamonds loops (10 unique motifs, Table 3), and 2 to 5-gene feedbacks loops. The nomenclature of the FFLs was based on (Mangan and Alon, 2003), the genes under consideration hold the position *Z* (regulated gene) in the motifs. FFLs were divided into two groups, coherent and incoherent. The coherence of a motif was defined by the product of all regulations of the motif: coherent if the product was positive, incoherent otherwise (Table 2). The nomenclature of diamonds

**Table 2:**
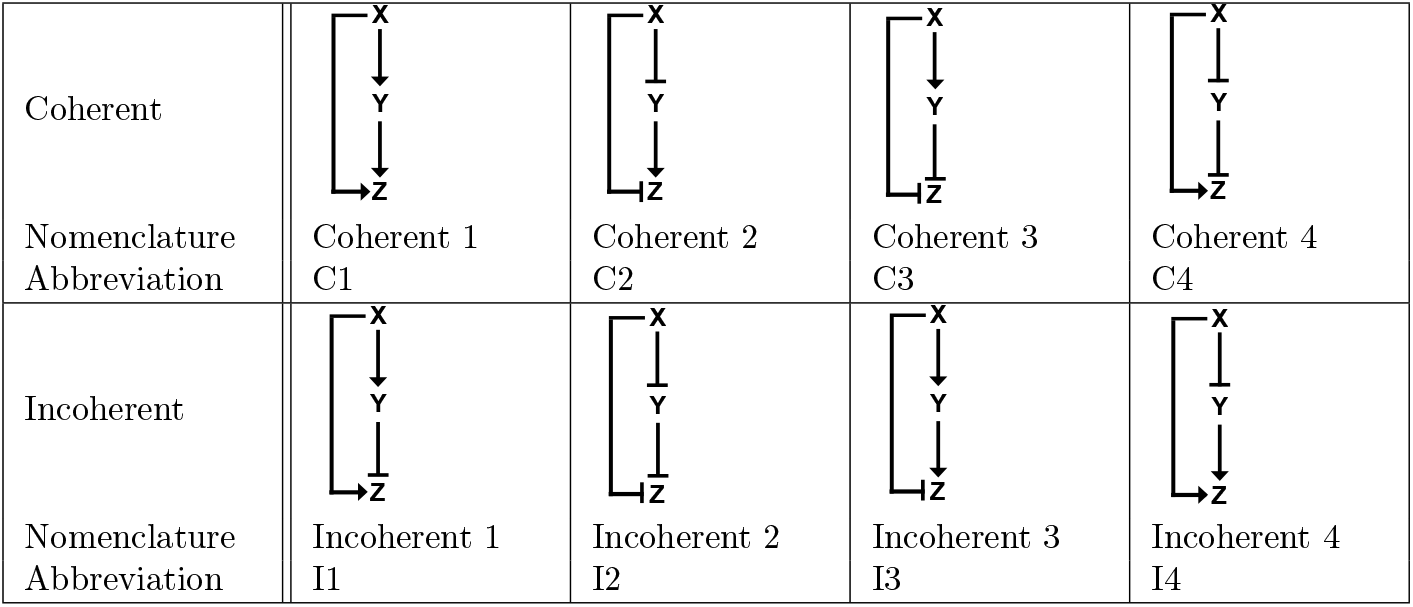
Three-gene feedforward loop nomenclature.

|  |  |  |  |  |
| --- | --- | --- | --- | --- |
| Coherent |  |  |  |  |
| Nomenclature Abbreviation | Coherent 1<br>C1 | Coherent 2<br>C2 | Coherent 3<br>C3 | Coherent 4<br>C4 |
| Incoherent |  |  |  |  |
| Nomenclature Abbreviation | Incoherent 1<br>I1 | Incoherent 2<br>I2 | Incoherent 3<br>I3 | Incoherent 4<br>I4 |

**Table 3:**
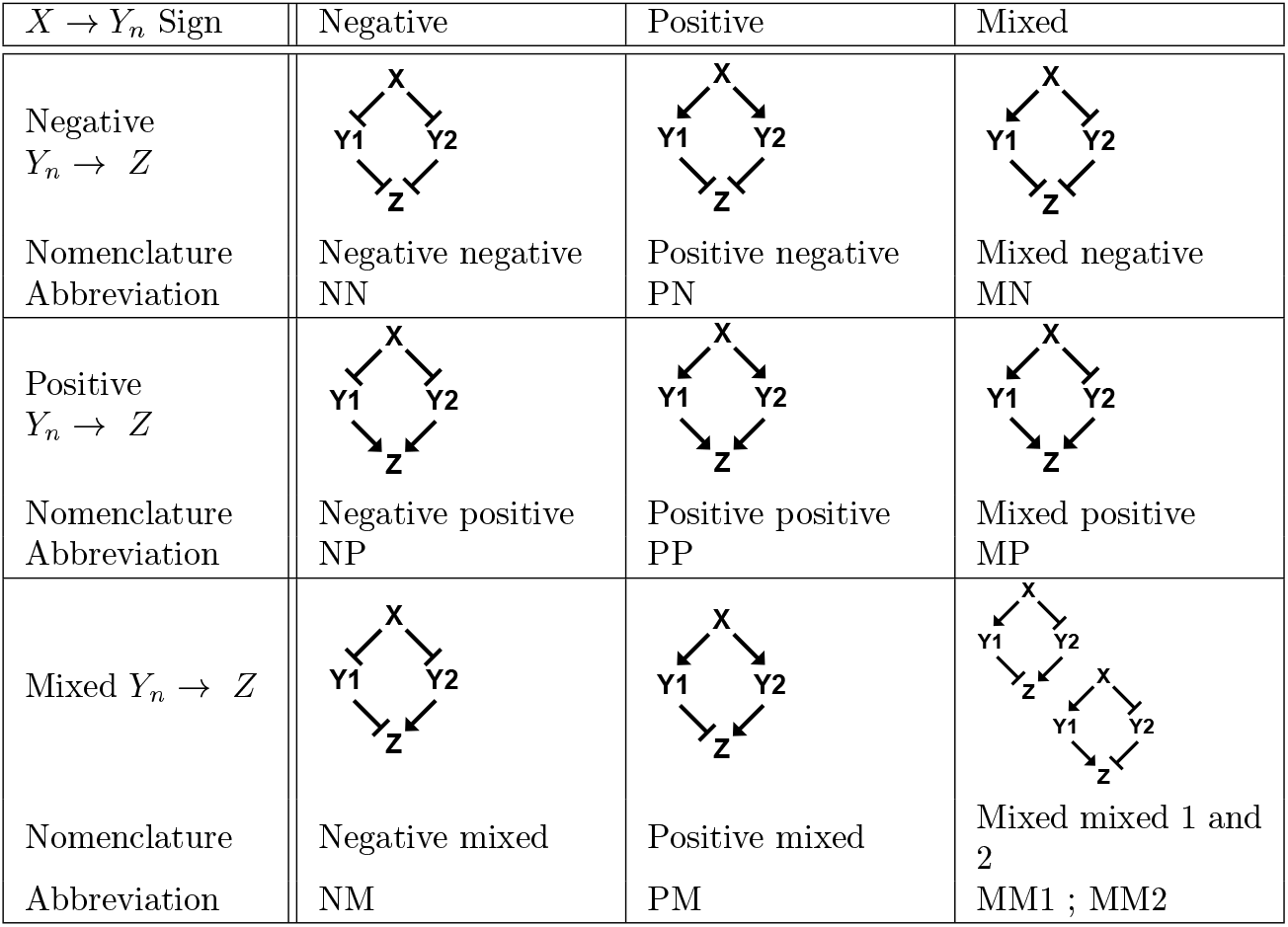
Diamond motifs nomenclature.

| $X \rightarrow Y_n$ Sign | Negative | Positive | Mixed |
| --- | --- | --- | --- |
| Negative<br>$Y_n \rightarrow Z$ | | | |
| Nomenclature<br>Abbreviation | Negative negative<br>NN | Positive negative<br>PN | Mixed negative<br>MN |
| Positive<br>$Y_n \rightarrow Z$ | | | |
| Nomenclature<br>Abbreviation | Negative positive<br>NP | Positive positive<br>PP | Mixed positive<br>MP |
| Mixed $Y_n \rightarrow Z$ | | | |
| Nomenclature<br>Abbreviation | Negative mixed<br>NM | Positive mixed<br>PM | Mixed mixed 1 and 2<br>MM1 ; MM2 |

was defined by the the signs of the *X* →*Y n* regulations (positive, negative, or mixed) for the first part of the name and the sign of *Y n* →*Z* for the second part (Table 3). Feedback loops were divided into subcategories by size, from 2 to 5 genes.

#### Control datasets

We considered two types of control: drifting simulation and randomized networks. In drifting simulations, populations of 200 individuals (i.e., smaller than in the main simulations) evolved for 20000 generations with a mutation rate of *µ* = 0.4 (larger than in the main simulations). “Shuffled” controls were obtained by randomly shuffling the values of non-null gene regulations. The number of regulations and loops toward genes were thus kept, but not the motifs.

### Availability and reproducibility

The simulation program was coded in C++, and is available at https://github.com/lerouzic/simevolv. The program was compiled and executed on the IFB-core cluster (https://www.france-bioinformatique.fr/en/ifb-core-cluster/). Simulation pipelines (in R and Bash) and analysis scripts (in R) are available at https://github.com/apetit8/Topoplast.

## Results

The *E. coli* regulatory network featured 3084 genes, including 326 regulators (graphic representation of the network in Suppl. Fig. 1). Out of the 4638 genes annotated in the reference annotation, 2144 (46%) genes were never reported as differentially expressed in our 11 reference studies, and were thus considered as “non-plastic”. In contrast, 1074 (23%) genes were differentially expressed in at least two studies, and therefore considered as “plastic” (Suppl. Table 1 and Suppl. Fig. 2). The 1420 (31%) genes reported only once as differentially expressed were excluded from the analysis.

Computer simulations were performed by coupling a dynamic gene network model and a classical population genetics framework. At the end of the simulations, genes were classified as plastic and non-plastic based on thresholds on the slope of the reaction norm; Because the sample size with simulation was virtually unlimited, we were more stringent and rejected 55% of the genes (mild plasticity). 73% of the remaining genes were clearly plastic, and 27% were clearly non-plastic.

### Reguation of plastic *vs.* non-plastic genes

We first considered the number of regulators of plastic and non-plastic genes, see Fig. 1A. Plastic genes were more regulated than non-plastic genes, both in *E. coli* and in simulations. In *E. coli*, plastic genes were regulated by 3.2 genes on average (7.2 in simulations), while non-plastic genes were regulated by 1.4 genes on average (4.2 in simulations, to be compared to the strict minimum of 1 regulator necessary to deviate from the constitutive expression). The mean number of regulators of *E. coli* plastic and non-plastic genes were found significantly different by a t-test (one-sample t-test, *t* = −11.3; *df* = 1480.3, *p* = 2· 10^*−16*^; statistical tests were conducted on *E. coli* data only, any difference being statistically supported in highly-replicated simulations). In absence of selection (drift simulations), genes that happened to evolve a plastic response were also characterized by a larger number of regulators, suggesting that plasticity tends to be passively induced by a larger number of regulators. In Fig. 1B, we looked at the number of feedforward, diamond, and feedback loops targeting genes. The number of loops targeting plastic genes was again larger than for non-plastic genes in both *E. coli* and in simulations, including in control simulations where mutations could accumulate freely with genetic drift. In drift simulations, there were more regulatory loops than in regular simulations for the same number of regulators, suggesting that stabilizing selection on gene expression tends to limit the number of regulatory loops.

**Figure 1.**
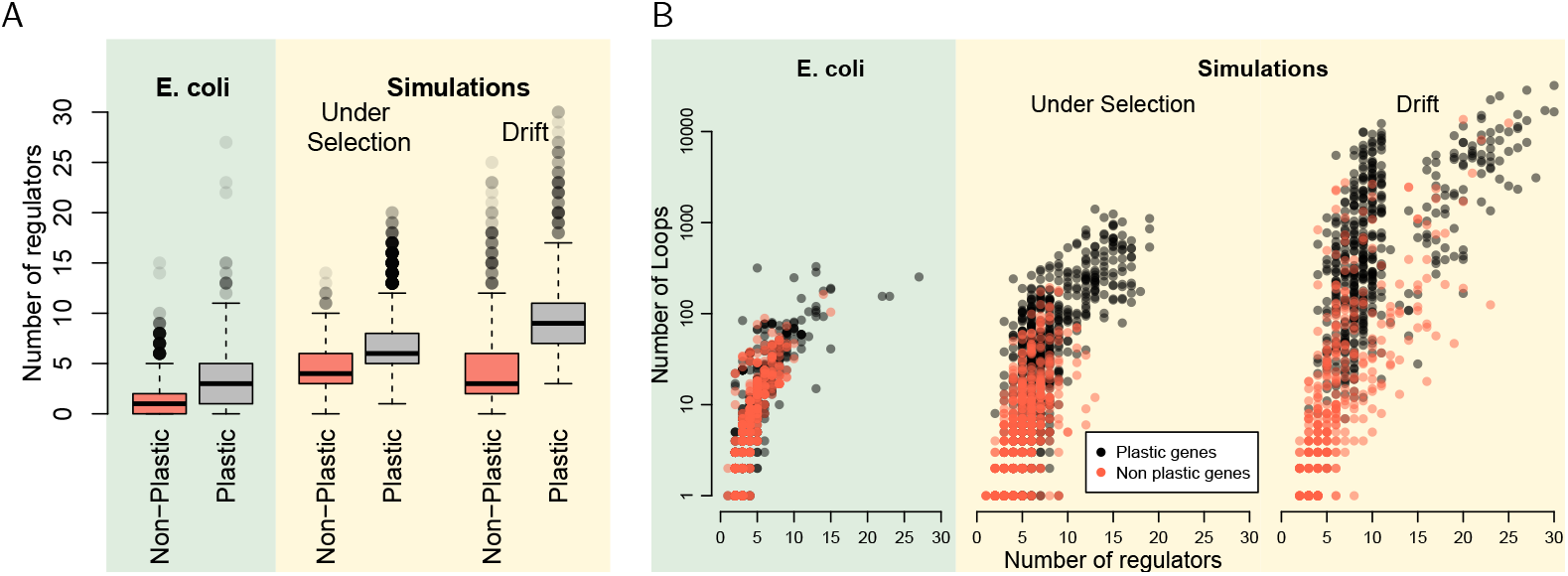
Regulation of plastic and non-plastic genes. A) Distribution of the number of regulations toward the genes. B) Number of feedforward, diamond, and feedback loops by which genes were regulated (only genes regulated by at least one loop are displayed). The discontinuity in the “drift” panel separates the transcription factors (which can get feedback from all the network genes, i.e. *>* 10 regulators) and the rest of the genes which can get regulated by 10 transcription factors at most. Simulation results have been downsampled to show the same number of data points in all panels.

### Regulatory loops

In *E. coli*, 64% of plastic genes (Fig. 2A), and 30% of non-plastic genes (Fig. 2B) were regulated by at least one of the loop categories considered (*i.e.* feedforward, feedback, and diamonds). The most common observed pattern for *E. coli* genes (plastic or not) was the regulation by both FFLs and DMDs, with a slight excess of FFLs compared to DMDs. Feedback loops (FBLs) involving 2 to 5 genes were very rare in *E. coli* (68 genes regulated by FBLs out of 3214). Overall, the proportions of genes regulated by FFLs, DMDs, FBLs, and any combination thereof were strikingly similar between plastic and non-plastic genes, and none of these loop categories appeared to be preferentially associated with plastic gene expression.

**Figure 2.**
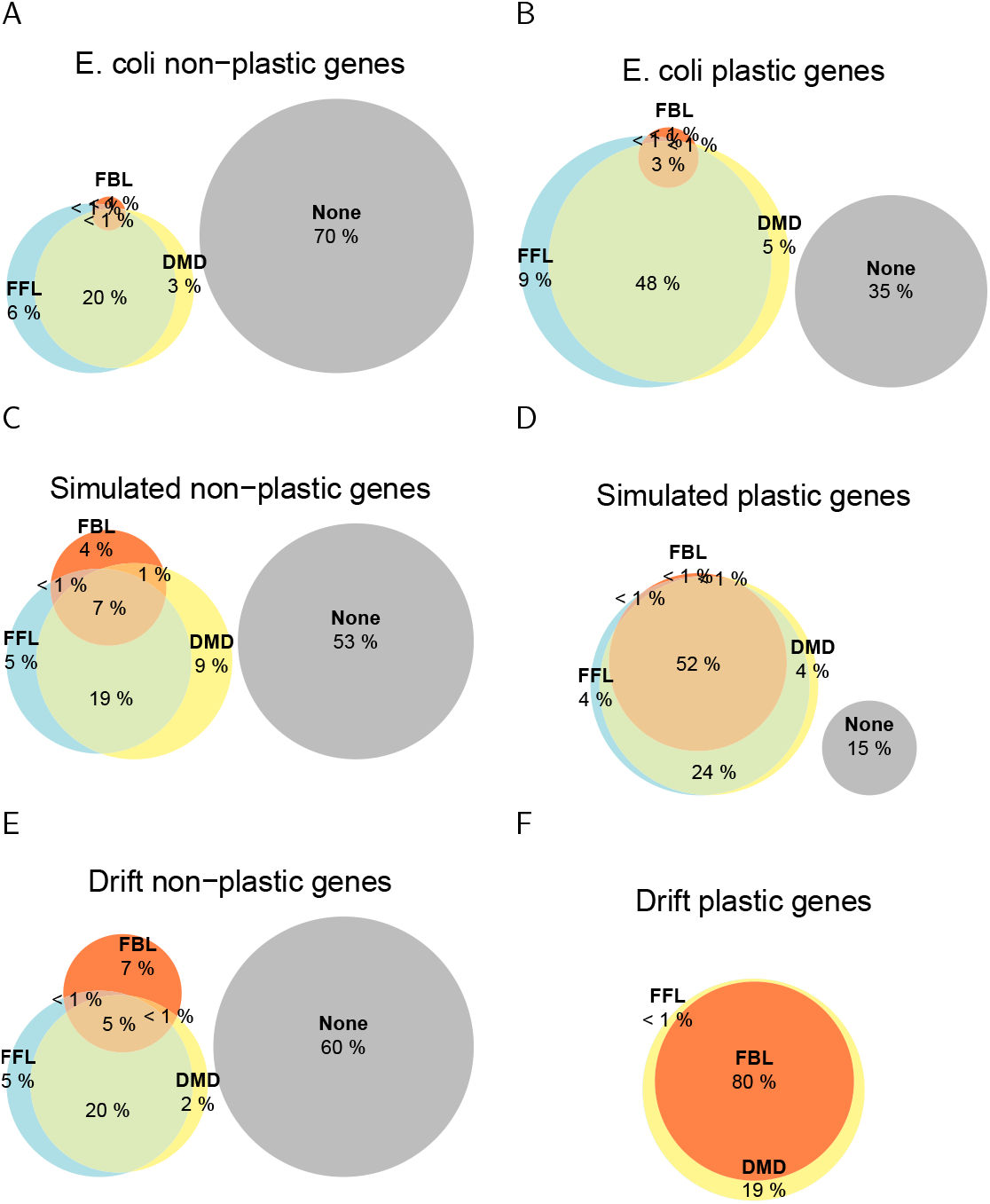
Genes classified by incoming regulation loops. A,C,E) Non-plastic genes in *E. coli*, simulations and drift. B,D,F) Plastic genes in *E. coli*, simulations, and drifting simulations. FFL: Feedforward loops, FBL: Feedback loops, DMD: Diamond loops. “None” stands for genes which regulators were unknown, or which were regulated by other motifs (no loops, or loops involving more or less genes than investigated here).

In simulated gene networks, plastic genes were also more frequently regulated by feedforward, feedback, or diamond loops than non-plastic genes. Most genes were targeted by several categories of loops, and the proportion of FFLs *vs.* DMDs was even (Fig. 2C and D). Overall, there was no clear difference in the relative proportion of FFL and DMD involved in the regulation of plastic *vs.* non-plastic genes. FBLs were common in simulated networks (although less frequent than DMD and FBLs), and were often associated with FFL and DMD. For non-plastic genes, the regulatory loops were similar between selection and drift simulation. However, this was not the case for plastic genes, which regulation involved more FBLs and virtually no FFLs when plasticity arose by drift. This points out a substantial effect of selection on the regulation of plastic genes; stabilizing selection on gene expression tends to homogenize the regulatory network topologies around plastic and non-plastic genes.

### Loop motifs occurrence

Loop categories remains a coarse-grained motif classification, as the precise regulatory function necessarily depends on the sign and the coherence of interactions. We have thus determined the motif percentage within FFL and DMD loops (8 motifs for FFLs, Table 2, and 10 motifs for DMD, Table 3), for both the *E. coli* dataset and for simulated networks. Feedback loops were too rare in *E. coli* networks (only 68), and thus not included in Fig. 3A and B.

**Figure 3.**
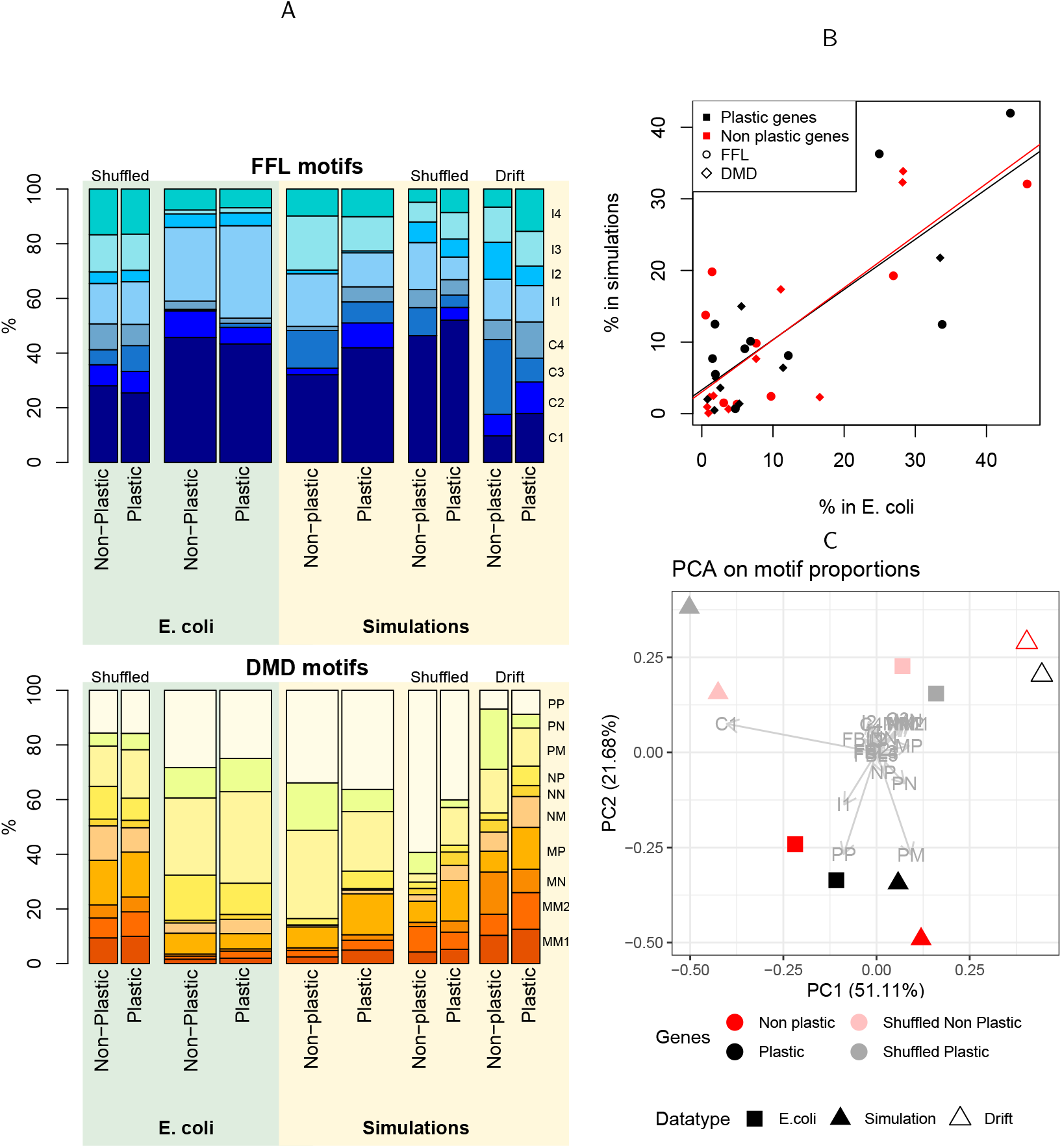
Loop motifs regulating plastic and non-plastic genes. A) Percentage of loop motifs within feedforward (top) and diamond (bottom) loop categories. Motif nomenclature corresponds to Tables 2 and 3. Motif percentages were normalized per gene (i.e., all genes have the same weight in the figure, regardless of the number of regulatory motifs that target them). Shuffled controls were obtained by randomly permuting all non-null interactions. Simulation runs in the absence of selection are indicated as “drift”. B) Correlation plot of DMD and FFL motif frequencies (shown in panel A) between simulation and *E. coli*. C) Principal component analysis on the percentage of loop motifs (FFL, DMD and FBL).

In both *E. coli* and simulated gene networks, the most frequent feedforward motif was C1 (coherent, two genes up-regulating), followed by I1 (incoherent, one gene up-regulating and the other downregulating) (Fig. 3A). Other motifs were found in similar proportions, with the exception of I3, frequent in simulations but almost absent in *E. coli*. For diamond loops, the most frequent motifs in both *E. coli* and simulations were PP (up-regulated by both genes) and PM (up-regulated by one gene, downregulated by the other one). The motifs frequency was similar among all the studies used to identify plastic genes (Suppl. Fig. 3). Motifs involving negative regulations (NN, NM, and NP) were more frequent in *E. coli* than in simulated networks; but overall, the motif distribution remained quite similar and the frequency of motifs found in *E. coli* and simulations was highly correlated, see Fig. 3B (plastic genes: r = 0.80; p-value = 5.9 · 10^*−5*^ ; non-plastic genes: r = 0.79 ; p-value = 1.0 · 10^*−4*^).

In contrast, the motif frequency distribution resulting from an evolutionary process occurring both in simulations and in *E. coli*, was substantially different from the randomly shuffled networks. In particular, the most frequent motifs (C1 and I3 for FFLs, PP and PM in DMDs) were enriched in evolved networks compared to the shuffled and drift controls. This enrichment was a consequence of selection for an optimal gene expression in simulations, and it is reasonable to postulate that the same explanation holds for *E. coli*.

While there was a clear contrast between control and selected networks, differences in motif composition between plastic and non-plastic genes for *E. coli* were non-significant. We performed Two-Proportion Z-tests for each motifs to determine which loop motifs differed between plastic and non-plastic genes. None of loop motifs were found to be significantly different in proportion between plastic and non-plastic genes after correction for multiple testing (Suppl. Table 2).

In a Principal Component Analysis on the relative proportion of each motifs of the three loop categories (Fig. 3C), PC1 (50% of the variance) was discriminating *E. coli* vs. simulations, while PC2 (22% of the variance) was associated with the contrast between selected and random networks. Plastic and non-plastic genes from the same conditions were found closer to each other than any other points, reinforcing that the contribution of plasticity to the distribution of motif frequency was minor. Moreover, *E. coli* and simulations were closer to each other than to their respective shuffled networks or drift.

### Discussion

Our objective was to identify the gene regulatory topologies involved in gene expression plasticity, from both empirical and evolutionary simulated regulatory networks. We found that plastic genes indeed had specific regulatory characteristics: genes which expression depends on the environment had more regulators, and were the target of more regulatory loops. The analysis of the frequency of the different regulatory loops and regulatory motifs showed that both real and simulated networks were not random, as a consequence of selection on gene expression. The difference between plastic and non-plastic genes seems to be related to the number of regulators, but we found no evidence of an association between plasticity and regulatory motif.

### Plastic genes have more regulators

We identified some regulatory differences between plastic genes and non-plastic genes in both real and simulated data. On average, plastic genes were regulated by more genes, and were the target of more regulatory loops (feedforward loops, diamond loops, and feedback loops). These results reinforce previous observations, which associated the number of regulators with the complexity of the gene expression dynamics (Promislow, 2005; Mochizuki, 2008; Bei *et al.*, 2024). Nevertheless, the distributions of the number of regulators and the number of regulatory loops largely overlap between plastic and non-plastic genes, both in simulated and in empirical networks. As a consequence, the number of regulators remains a poor predictor of gene expression plasticity.

An obvious advantage from simulations to complement the analysis of empirical network lies in the possibility to go beyond association statistics. We showed for instance that plasticity was statistically associated with a higher number of regulators, even in the drift simulations. We suspect here that increasing the number of regulators increases the probability to connect the gene to a subnetwork influenced by the environmental factor, thus enriching the proportion of plastic genes among highly-connected genes. We propose the following scenario for the changes in the regulatory networks associated with the evolution of plasticity: selection acting on plastic gene expression directly promotes the recruitment of *cis*-regulators, which indirectly increases the number of regulatory loops. Thus, the reported association between some regulatory loops like FFL with plasticity may only be a consequence of the higher number of regulation towards plastic genes.

We also obtained interesting contrasts between selected and drift simulations, since selection tends to (i) decrease the number of loops for a given number of regulators, and (ii) homogenize the distribution of loops and loop motifs between plastic and non-plastic genes. Selection for gene expression control thus implies the elimination of supernumerary and/or inadequate loops and loop motifs, and the enrichment in motifs such as C1, I1, PP, and PM. C1 and I1 have for instance been predicted to be efficient at buffering expression noise (Ghosh *et al.*, 2005; Osella *et al.*, 2011; Reeves, 2019; Chakravarty and Csikász-Nagy, 2021). Strikingly, this selection-related enrichment makes plastic and non-plastic gene regulation more similar than what could be observed without selection. Therefore, the regulation of plastic vs. non-plastic genes could theoretically be different (as we observed in the drift simulations for instance), but they happen to be similar because the regulatory structure was overwhelmed by the necessity to control for gene expression (for a fixed or an environmentally-driven optimal value, indifferently).

The *E. coli* motif distribution was consistent with previous work (Mangan *et al.*, 2006; Cooper *et al.*, 2008), who also identified C1 and I1 as the most represented motifs. Up-regulating motifs (including C1 for feed-forward loops or PP for diamond loops) were over-represented in *E. coli* compared to randomly-permuted networks, suggesting some specific preference for such motifs. However the high frequency of up-regulating motifs (including C1 for feed-forward loops or PP for diamond loops) could also partially be a statistical consequence of the uneven ratio between up- and down-regulation within the network. “Pure” up-regulating motifs C1 and PP were indeed rare in simulations evolving by drift only, featured by an equal number of up- and down-regulation interactions. Yet, arguments related to up/down-regulation ratios are fragile, as it is difficult to discard that the uneven balance could be a consequence, rather than a cause, of an evolutionary bias towards specific motifs. In our simulations, the gene network dynamics was deterministic, so that it was impossible to disentangle the control of the equilibrium expression from the control of random perturbations (noise). Alternative theoretical settings as proposed in Draghi and Whitlock, 2015; Schmutzer and Wagner, 2020 could be considered in the future to address this interesting issue.

### Theoretical and empirical results are converging

Systems biology is rather convincing at describing general patterns in biological networks, based on measurement tools derived from graph theory (Mason and Verwoerd, 2007; Pavlopoulos *et al.*, 2011). For instance, it is often admitted that biological networks tend to be scale-free (Babu *et al.*, 2004), relatively scarce (Espinosa-Soto, 2018), and modular (Wagner *et al.*, 2007). Yet, there is a surprising lack of theory from which such patterns could be interpreted. Typically, the extent to which network evolution is driven by mutation bias, genetic drift, adaptation, or indirect selection remains widely unknown (Tsuda and Kawata, 2010; Halfon, 2017; Petit *et al.*, 2023). Here, we show how a simplified gene network evolutionary model, generally exploited to address genetic architecture-related questions, can nourish the field of systems biology and predict empirical patterns. Our simulations could retrieve, both qualitatively and quantitatively, the larger number of regulatory genes for plastic genes, motif enrichment and depletion in selected *vs.* randomized networks, the most abundant motifs, and the relationship between the number of regulation and number of motifs. This validates the capacity of such simple models in which gene regulations are encoded in an interaction matrix (Wagner, 1994; Fierst, 2011), to contribute to the theoretical corpus on gene network evolution, in complement to the more cumbersome physiologically-realistic models (Xiong *et al.*, 2019).

Nevertheless, we noticed some differences between empirical and simulations results, that might be worthwhile to explore. First, the most obvious qualitative difference between both approaches was the ∼10-fold decrease in the frequency of feedback loops in *E. coli* regulation network compared to simulations. Although feedback regulation exists in *E. coli* and is known to be involved into noise reduction or bistability (Smits *et al.*, 2006), feedback loops happen to be outnumbered by feedforward loops in the Ecocyc transcriptional dataset and in pathway databases in general (Karp, 2001; Kumar *et al.*, 2018). A possible explanation for this discrepancy lies in the fact that most feedback mechanisms in prokaryotes rely on autoregulation (in *E. coli*, 50% of TFs self-regulate, Rosenfeld *et al.*, 2002), while the minimum size for a motif in our analysis was set to 2 genes. Secondly, a more subtle discrepancy between theoretical and empirical networks was that simulations predicted a small but systematic difference between motif frequencies in plastic vs. non-plastic genes, which was not statistically supported in *E. coli*. Insufficient statistical power could partly explain this observation, but more generally, the classification of *E. coli* genes into plastic and non-plastic categories remained coarse-grained, and may not match the same definition of plasticity; simulations define plastic genes based on the slope of the reaction norm to a single environmental factor, while in *E. coli* we defined plastic genes based on their differential expression to different environmental treatments, which might not reflect exactly the same mechanisms. The observed convergence between theoretical and empirical observations in spite of these different measurements of plasticity strengthens our conclusions.

In our model, the ratio of up-vs. down-regulation interactions was not a parameter, but rather the outcome of the evolutionary process. As a result, this rate could not be easily controlled and ended up to differ from the observed network database (78% of up-regulation in simulations vs. 61% in *E. coli*). This difference limits the robustness of quantitative conclusions, as it conditions the expected proportions of the loop motifs (differences in the “shuffled” control in simulations vs. *E. coli*). We are therefore confident that the striking correlation between motif frequencies in simulations vs. *E. coli* reflects the capacity of the model to simulate realistic evolutionary trajectories, but we cannot be certain that this convergence shoud be attributed to analogous selection pressures since motif frequencies had to evolve from a different baseline.

### Conclusion

If not the network topology, which regulatory features could reveal whether the expression of a specific gene has been selected for robustness, stability, cyclic expression, or sensitivity to an environmental or physiological input? To what extent is it possible to predict gene expression profiles from pathway databases (Fang *et al.*, 2017; Mercatelli *et al.*, 2020; Escorcia-Rodríguez *et al.*, 2023)? Three major hypotheses can be proposed: (i) the information about network topologies involved in plasticity is present in existing databases, but they were not caught by known descriptors of network topologies and properties (loops, motifs, or graph metrics). For instance, it might depend on a more general regulatory context (Lipshtat *et al.*, 2008; Jiang and Hao, 2021), or on complex combinations of several intertwined network topologies (Chakravarty and Csikász-Nagy, 2021). Alternatively, (ii) regulation databases remain the correct approach, but existing databases are not precise nor exhaustive enough. Typically, presence/absence data for regulatory interactions might be insufficient, and quantitative estimation of interaction strengths (weighted edges in the network graph (Freyre-González *et al.*, 2022)) could be necessary as suggested by others (Ingram *et al.*, 2006; Panovska-Griffiths *et al.*, 2013; Kim and Wysocka, 2023). The last hypothesis (iii) presumes that regulation might be too complex to be abstracted in regulatory networks, which omit important epigenetic and cellular mechanisms, including the spatial organization of the genome, methylation, phosphorylation, dimerization (Klumpe *et al.*, 2023), protein conformation, or “missing regulation” from intergenic sequences (Connally *et al.*, 2022). The next challenge for the systems biology community may therefore not be to generate more data, but to identify which data is relevant to understand the evolutionary properties of biological systems, a task for which theoretical approaches might bring invaluable insights.

## Aknowledgments

Simulations were performed on the Core Cluster of the Institut Français de Bioinformatique (IFB) (ANR-11-INBS-0013). AP was supported by the doctorate school SDSV (ED 577, Université Paris-Saclay). The authors acknowledge the support of the French Agence Nationale de la Recherche (ANR), under grant ANR-22-CE02-0026-01 (project EVOPLANET). The authors declare no conflict of interest.

