## Supplemental tab and figures for "Gene Expression Plasticity Is Associated with Regulatory Complexity but Not with Specific Network Motifs"

| | Tested environmental variable | Tucker 2002 | Maurer 2004 | Bhatia 2022 | Feugas 2016 | Caglar 2017 | Wang 2009 | Ng 2018 | Bergholz 2009 | White-Ziegler 2007 | Kim 2020 | Durfee 2008 |
|---|---|---|---|---|---|---|---|---|---|---|---|---|
| Tucker 2002 | pH | 30 | 0 | 13 | 2 | 16 | 4 | 4 | 0 | 0 | 4 | 4 |
| Maurer 2004 | pH | | 667 | 26 | 86 | 363 | 37 | 13 | 28 | 31 | 78 | 158 |
| Bhatia 2022 | Metabolic stress | | | 147 | 27 | 76 | 15 | 2 | 9 | 6 | 18 | 19 |
| Feugas 2016 | Medium of growth | | | | 390 | 207 | 25 | 5 | 10 | 28 | 33 | 60 |
| Caglar 2017 | Sources of C and Mg | | | | | 1694 | 76 | 29 | 49 | 60 | 156 | 270 |
| Wang 2009 | Oxidative stress | | | | | | 127 | 3 | 8 | 14 | 22 | 23 |
| Ng 2018 | Aerosolization | | | | | | | 59 | 5 | 6 | 9 | 8 |
| Bergholz 2009 | Exposure to apple juice | | | | | | | | 112 | 1 | 9 | 23 |
| White-Ziegler 2007 | Temperature | | | | | | | | | 120 | 17 | 7 |
| Kim 2020 | Temperature | | | | | | | | | | 217 | 41 |
| Durfee 2008 | amino acid starvation | | | | | | | | | | | 513 |

Supplementary Table 1: Table of the number of reported environmentally differentially expressed gene in each study (diagonal) and number of common plastic genes between pairs of studies (off-diagonal).

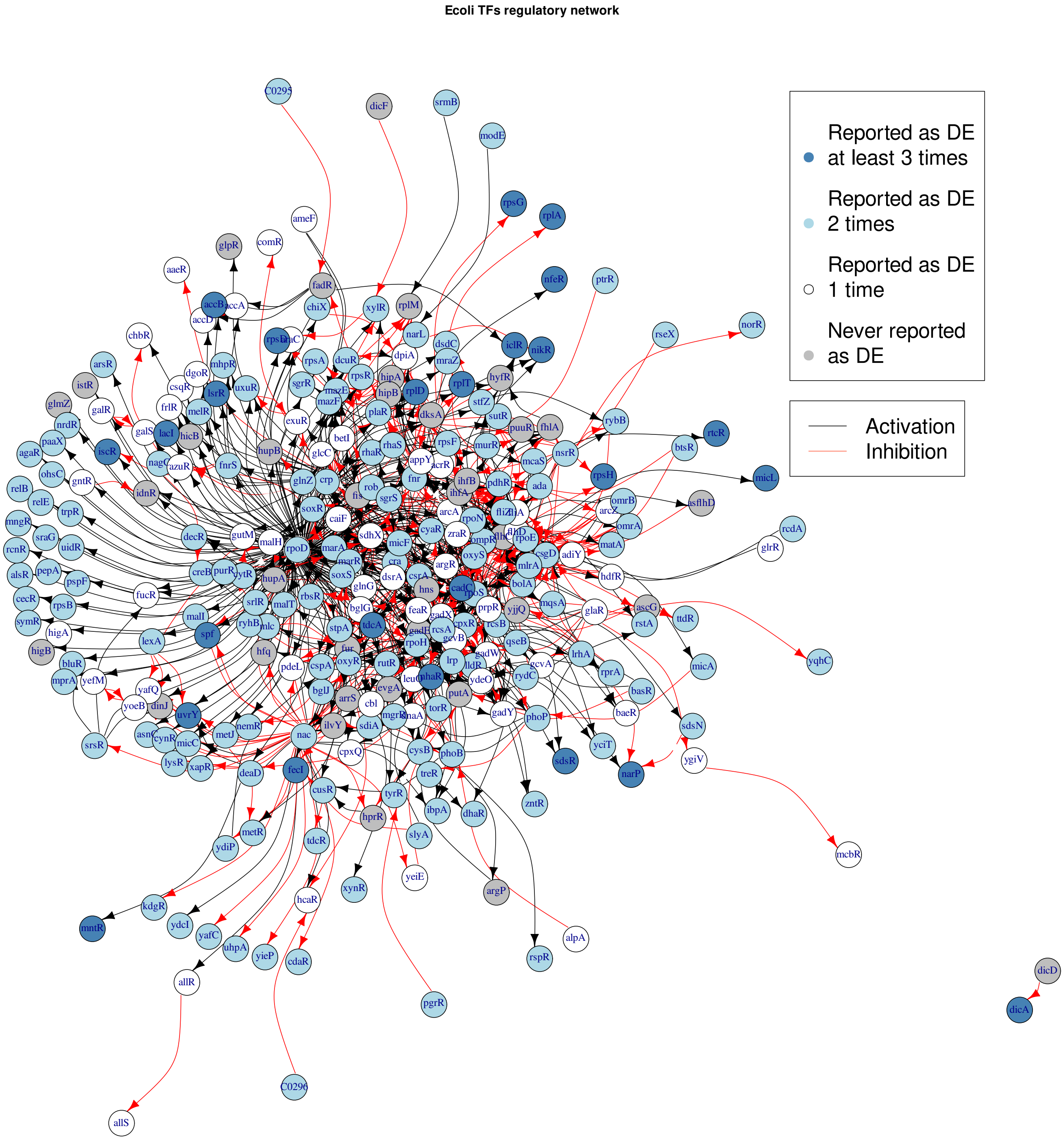


Supplementary Figure 1: Representation of gene regulations between the transcription factors of *E. coli*. Self-regulations are not displayed. Data from Ecocyc.

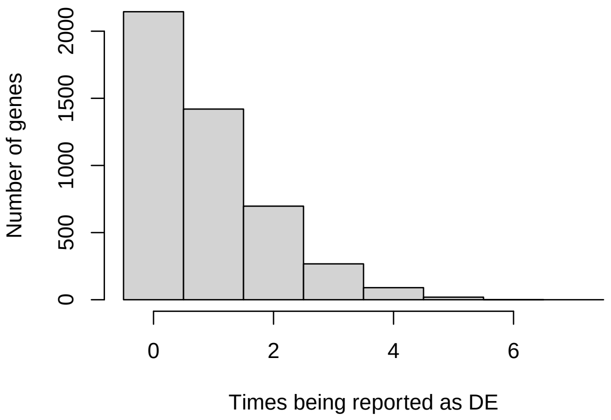


Supplementary Figure 2: Distribution of the number of times genes were reported as differentially expressed in response to an environmental change in the considered studies.

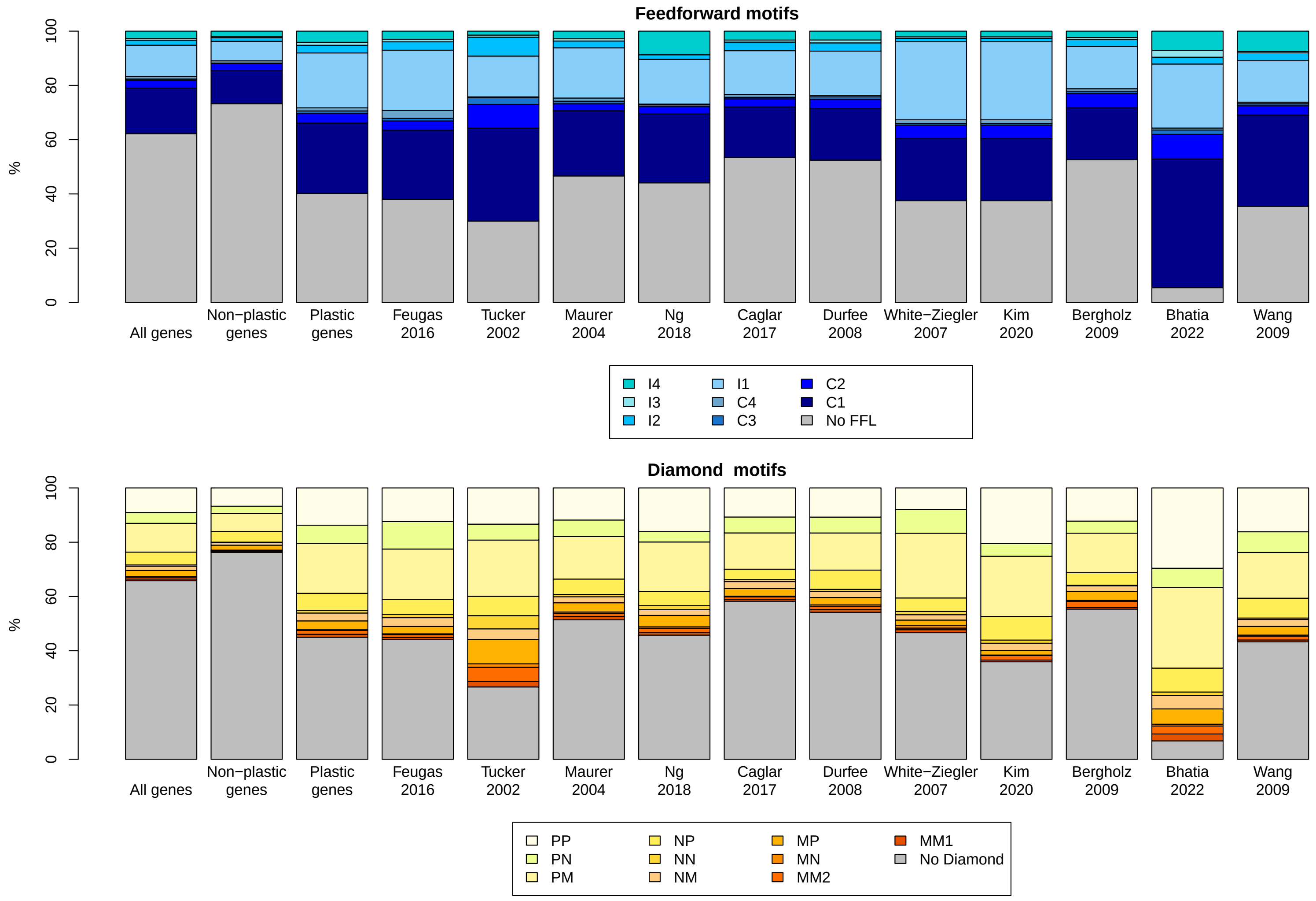


Supplementary Figure 3: Frequency of motifs regulating differentially-expressed genes in *E. coli*.

| Motif | Non-plastic genes | Plastic genes | P-value | Q-value |
|---|---|---|---|---|
| FFL | | | | |
| C1 | 261.9 | 278.8 | 0.44 | 0.81 |
| C2 | 55.7 | 38.8 | 0.02 | 0.16 |
| C3 | 3.1 | 9.7 | 0.16 | 0.61 |
| C4 | 17.6 | 12.3 | 0.26 | 0.63 |
| I1 | 154.2 | 217.2 | 0.01 | 0.16 |
| I2 | 28.3 | 30.3 | 0.96 | 1 |
| I3 | 8.3 | 11.9 | 0.74 | 0.99 |
| I4 | 43.8 | 44.1 | 0.67 | 0.98 |
| Number of genes | 573 | 643 | NA | NA |
| DMD | | | | |
| PP | 144 | 147.4 | 0.24 | 0.63 |
| PM | 143.6 | 197.9 | 0.07 | 0.37 |
| PN | 56.5 | 71.8 | 0.65 | 0.98 |
| NP | 84.4 | 67.5 | 0.02 | 0.16 |
| NM | 19.1 | 31.1 | 0.29 | 0.64 |
| NN | 4.8 | 10.5 | 0.36 | 0.71 |
| MP | 38.8 | 32.9 | 0.21 | 0.63 |
| MM2 | 5.7 | 15.4 | 0.11 | 0.50 |
| MM1 | 8.3 | 11.8 | 0.82 | 1 |
| MN | 3.9 | 4.7 | 1 | 1 |
| Number of genes | 509 | 591 | NA | NA |
| FBL | | | | |
| FBL2 | 7.3 | 7.1 | 0.56 | 0.94 |
| FBL3 | 0.9 | 2.7 | 0.94 | 1 |
| FBL4 | 5.2 | 7.2 | 1 | 1 |
| FBL5 | 11.7 | 20.1 | 0.76 | 0.99 |
| Number of genes | 25 | 37 | NA | NA |

Supplementary Table 2: The proportion of motif-regulated genes that are subject to regulation by the different motifs. Two-Proportion Z-tests were conducted for each motif, and q-values were obtained by the false-discovery-rate method.
